# HID-1 controls cargo sorting and dense core formation by influencing *trans*-Golgi network acidification in neuroendocrine cells

**DOI:** 10.1101/156794

**Authors:** Blake H. Hummer, Noah F. de Leeuw, Bethany Hosford, Christian Burns, Matthew S. Joens, James A.J. Fitzpatrick, Cedric S. Asensio

## Abstract

Large dense core vesicles (LDCVs) mediate the regulated release of neuropeptides and peptide hormones. They form at the *trans*-Golgi network (TGN) where their soluble content aggregates to form a dense core, but the mechanisms controlling biogenesis are still not completely understood. Recent studies have implicated the peripheral membrane protein HID-1 in neuropeptide sorting and insulin secretion. Using CRISPR/Cas9, we generated HID-1 KO rat neuroendocrine cells, and show that the absence of HID-1 results in specific defects in peptide hormone and monoamine storage and regulated secretion. Loss of HID-1 causes a reduction in the number of LDCVs and affects their morphology and biochemical properties due to impaired cargo sorting and dense core formation. HID-1 KO cells also exhibit defects in TGN acidification together with mislocalization of the Golgi-enriched vacuolar H^+^-ATPase subunit isoform a2. We propose that HID-1 influences early steps in LDCV formation by controlling dense core formation at the TGN.

## Introduction

The ability to control the secretion of proteins regulates many aspects of biology, including physiology, development and behavior. The regulated secretion of peptide hormones, such as insulin and glucagon, is essential for glucose homeostasis, and defects in their secretion lead to serious metabolic deficits. The regulated release of many neuropeptides by the hypothalamus contributes to the modulation of brain function and controls a variety of physiological behaviors ranging from feeding to sleep and reproduction (Argiolas and Melis, 2013, Richter et al., 2014, Sohn et al., 2013).

In addition to the constitutive secretory pathway enabling the immediate release of newly synthesized proteins, specialized secretory cells such as neurons and endocrine cells also express a regulated secretory pathway (RSP). This pathway enables them to store a subset of secretory proteins (e.g., peptide hormones, neuropeptides) into vesicles that accumulate intracellularly. These vesicles are called secretory granules or large dense core vesicles (LDCVs), and their exocytosis can be triggered by an extracellular, physiological stimulus, leading to an increase in cytosolic Ca^2+^ concentration. The mechanisms controlling this regulated exocytosis depend on the presence of specific membrane proteins present both at the plasma membrane and on LDCVs, including synaptotagmins that act as Ca^2+^ sensors for vesicle fusion and release (Sudhof, 2012). Despite the significance of peptide hormones and neuropeptides for physiology and many human diseases, the cellular and molecular mechanisms controlling the biogenesis of LDCVs, which determines this molecular composition, still remains poorly understood.

Analysis of budding using metabolic labeling in rat neuroendocrine PC12 cells has demonstrated that LDCVs form at the *trans*-Golgi network (TGN) where sorting of soluble regulated secretory proteins from constitutively secreted proteins occurs (Tooze and Huttner, 1990). LDCVs contain large amount of “granulogenic” proteins, such as members of the granin family, that aggregate to form a dense core under the specific pH and redox conditions of the TGN (Chanat and Huttner, 1991, Gerdes et al., 1989). It has been proposed that these lumenal interactions represent a driving force in LDCV formation (Arvan and Halban, 2004), with proteins destined for other organelles being removed from nascent LDCVs after budding. This process of maturation can last up to several hours after budding and depends on clathrin as well as the adaptor proteins AP-1, GGAs and PACS (Dittie et al., 1996, Dittie et al., 1997, Kakhlon et al., 2006). However, even immature LDCVs can undergo regulated release in rat neuroendocrine PC12 cells (Tooze et al., 1991), suggesting that LDCV maturation might not be a strict functional requirement for all secretory cells.

Recent studies have revealed the contribution to neuroendocrine secretion of several other cytosolic factors controlling various aspects of LDCV formation and maturation. For example, the EARP complex and its interactor EIPR-1 (Topalidou et al., 2016), as well as the adaptor protein AP-3 (Asensio et al., 2010, Sirkis et al., 2013) and its putative coat VPS41 (Asensio et al., 2013) have been involved in cargo sorting to LDCVs, whereas the BAR (Bin-Amphiphysin-Rev) domain protein Arfaptin-1 has been proposed to influence the size of insulin granules by regulating the timing of Arf1-mediated scission at the TGN (Gehart et al., 2012). The BAR domain proteins (Pick1 and ICA69) are thought to bind to immature LDCVs to regulate their number and size (Cao et al., 2013, Holst et al., 2013). Finally, several Rab proteins (Rab-2, Rab-5 and Rab-10), as well as Rab-2 effectors (Ailion et al., 2014, Edwards et al., 2009, Hannemann et al., 2012, Pinheiro et al., 2014, Sasidharan et al., 2012, Sumakovic et al., 2009) are involved in LDCV maturation.

A forward genetic screen in the model organism *C. Elegans* has previously identified HID-1 as a factor implicated in neuropeptide sorting and secretion (Mesa et al., 2011). HID-1 null worms display reduced levels of LDCV soluble cargo and impaired neurosecretion (Mesa et al., 2011, Yu et al., 2011), and conditional knockout (KO) mice lacking HID-1 in beta-cells of the pancreas display a defect in insulin secretion (Du et al., 2016), suggesting that this factor might be significant for glucose homeostasis. Interestingly, HID-1 is a peripheral membrane protein associated with the Golgi/TGN and its expression seems restricted to specialized secretory cells (Wang et al., 2011), suggesting that it might directly contribute to LDCV biogenesis. Here we test this hypothesis and show that HID-1 promotes mammalian neuroendocrine secretion by influencing LDCV cargo sorting and dense core formation. We further demonstrate that HID-1 is required for TGN acidification and propose that it controls an early step in LDCV formation at the TGN.

## Results

### HID-1 is required for LDCV soluble cargo storage and secretion

To investigate a potential role for HID-1 in mammalian RSP, we generated HID-1 knockout PC12 cells using genome editing. Specifically, we relied on homologous recombination repair after cleavage by Cas9 to knock-in a fluorescent protein (tdTomato), while effectively knocking-out HID-1. Our repair template was designed in such a way that the fluorescent reporter should not be expressed without proper, in-frame recombination (**Figure S1**). Although this approach does not guarantee bi-allelic homologous recombination, we reasoned that, as the efficiency of Cas9-mediated cleavage is much higher than homologous recombination repair, cells with monoallelic recombination would have a high probability of exhibiting indels on the other allele due to non-homology end joining. After transfecting PC12 cells with Cas9, gRNA and our repair template, we sorted cells positive for tdTomato by flow-cytometry. After three consecutive rounds of cell sorting, the proportion of positive cells shifted from an initial 0.2% to greater than 90%. To confirm that HID-1 was indeed deleted from our cell population, we analyzed our cells by immunofluorescence testing several commercial antibodies against HID-1, and identified one mouse monoclonal antibody (see Material and Methods) displaying a specific signal by immunofluorescence (**Figure 1A**). Consistent with our cell sorting enrichment data, we observed that HID-1 was absent from greater than 90% of our cells by immunofluorescence. This genome-editing approach offers the advantage of relying on transient expression of Cas9 and gRNAs, which should limit off-target effects.

**Figure 1.**
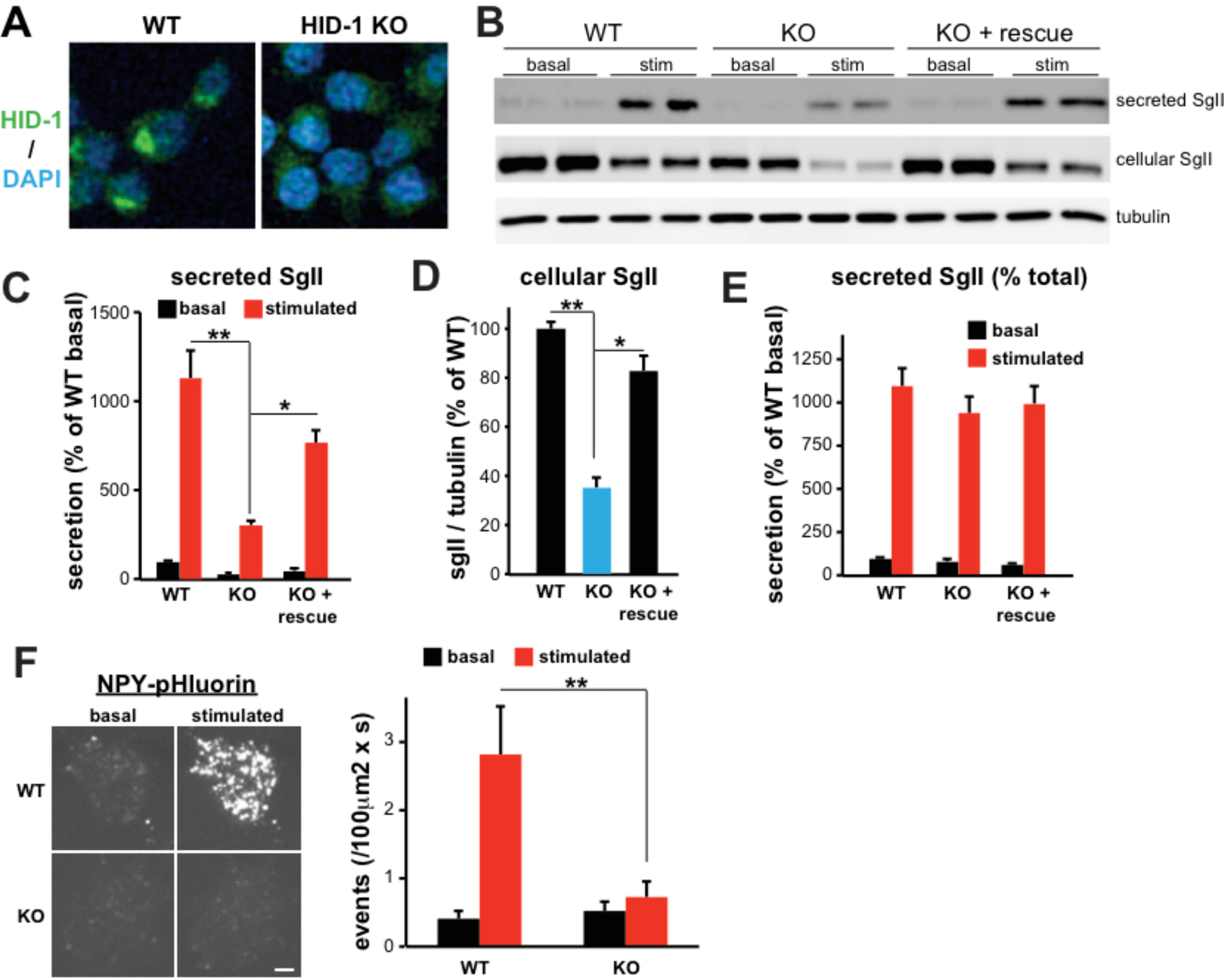
HID-1 is required for SgII storage and secretion from PC12 cells. (**A**) WT and HID-1 KO PC12 cells were stained with a mouse monoclonal antibody to HID-1, followed by an anti-mouse antibody conjugated to Alexa Fluor 488 and mounted with Fluoromount containing DAPI. Representative confocal micrographs show the absence of HID-1 staining in HID-1 KO PC12 cells. (B) PC12 cells (WT, HID-1 KO or HID-1 KO transduced with HID-1-HA lentivirus) were washed and incubated for 30 min in Tyrode’s solution containing 2.5 mM K^+^ (basal) or 90 mM K^+^ (stimulated). Cellular and secreted secretogranin II (SgII) were measured by quantitative fluorescent immunoblotting (**B**), with the secreted SgII normalized to tubulin (**C**) or total SgII (**E**) and expressed as percent of basal secretion in the control, and the basal cellular SgII normalized to tubulin (**D**). Multiple comparison statistical analysis was performed by one-way ANOVA followed by posthoc Tukey test *, p < 0.05; **, p < 0.01 relative to KO (n=4-6). No statistical difference was observed between WT and KO + rescue. The bar graphs indicate mean ± s.e.m. (**F**) WT and HID-1 KO PC12 cells were transfected with NPY-pHluorin, then imaged live by spinning disk confocal microscopy for 15 seconds in basal Tyrode’s solution. Regulated exocytosis was triggered by the addition of an equal volume of 90mM K^+^ Tyrode’s solution (45 mM K^+^ final) and imaged for an additional 30 seconds. Images show representative maximum intensity time projections of 150 basal and stimulated frames. At the end of the experiment, cells were imaged in Tyrode’s solution containing 50 mM NH_4_Cl, pH 7.4 to reveal total NPY-pHluorin fluorescence by alkalinization and identify transfected cells. Scale bar indicates 5 μm. Bar graph shows the number of exocytotic events per second normalized to cell surface area. Multiple comparison statistical analysis was performed by one-way ANOVA followed by posthoc Tukey test **, p < 0.01 relative to stimulated exocytosis from WT (n=11-15 cells from two independent experiments).

Loss of HID-1 significantly reduced basal cellular levels of the LDCV soluble marker secretogranin II (SgII) to ~40% of that observed in control cells by fluorescent western blot, and impaired the stimulation of SgII release by depolarization (**Figure 1B-D**). However, we found no significant reduction after normalizing secreted SgII values to total SgII levels, suggesting that there is no impairment in regulated exocytosis *per se* and that the decrease in SgII content is the major contributor to the defect in release (**Figure 1E**). We observed a similar phenotype in a second HID-1 KO PC12 cell line generated using an independent gRNA (**Figure S2**). Furthermore, we observed no change in SgII mRNA levels by qPCR (**Figure S3**) and found a similar storage and secretion defect of a transfected, exogenous soluble LDCV marker (ANF-GFP), whose expression is under the control of a strong CMV promoter, suggesting that the decrease in cellular content is unlikely to be transcriptional (**Figure S4**). Importantly, lentivirus-mediated expression of full-length HID-1 bearing a C-terminal HA-tag (HID-1-HA) in HID-1 KO cells restored SgII to WT levels and rescued the secretion phenotype (**Figure 1B-E**), thus ruling out Cas9 off-target effects.

To further assess secretion of LDCV soluble cargo, we transfected WT and HID-1 KO PC12 cells with NPY fused to the superecliptic pHluorin (NPY-pHluorin), which has been shown to undergo regulated exocytosis (Kogel et al., 2010, Sirkis et al., 2013), and monitored individual exocytotic events by live-imaging using spinning disk confocal microscopy (**Figure 1F, Movies S1,2**). As expected, WT PC12 cells displayed very few events under basal conditions and responded to depolarization (~7 fold over basal). The number of basal events observed in HID-1 KO cells remained low and unchanged, but we observed a striking reduction in the amount of exocytotic events in response to stimulation in these cells (**Figure 1F).** Altogether, our results suggest that the absence of HID-1 reduces storage and secretion of LDCV cargoes.

### HID-1 does not influence the endolysosomal or constitutive secretory pathway

To rule out that the impact of HID-1 KO on the RSP was due to indirect effects on constitutive secretion or endocytosis, we analyzed secreted fractions obtained from WT and HID-1 KO PC12 cells under non-stimulatory conditions by Coomassie (data not shown) or by silver staining after SDS-PAGE, and observed no obvious difference in the general pattern of constitutively secreted proteins (**Figure 2A**). Consistent with this, direct quantification of the secretion of emerald GFP fused to a signal sequence (ssGFP) revealed no change in release of this exogenous marker of the constitutive secretory pathway (**Figure 2B**).

**Figure 2.**
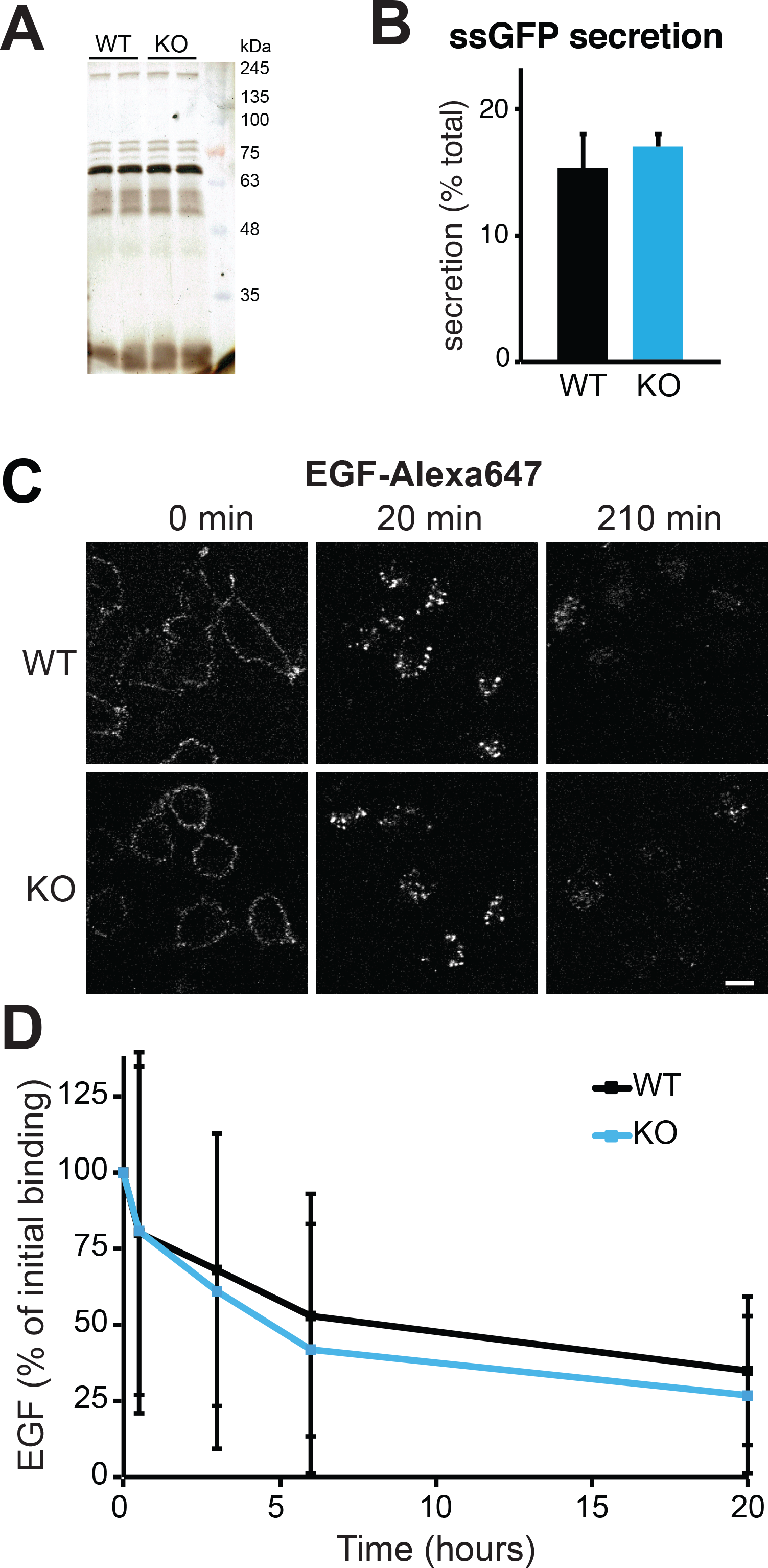
The absence of HID-1 does not impair the endolysosomal and constitutive secretory pathways. (**A**) Constitutive secreted fractions from unstimulated cells were analyzed by silver-staining after SDS-PAGE. (**B**) WT and HID-1 KO PC12 cells were transiently transfected with ssGFP, washed and incubated for 30min in basal Tyrode’s solution. Cellular and secreted ssGFP were measured using a plate reader, with the secreted signal normalized to cellular content (n=4). The bar graphs indicate mean ± s.e.m. (**C-D**) WT and HID-1 KO PC12 cells were incubated on ice with Alexa647 labeled EGF, and chased in complete media for the indicated time before fixation. Cells were then analyzed by confocal microscopy (**C**). The scale bar indicates 5 μm. (**D**) Alternatively, fluorescence was quantified by flow cytometry. Error bars indicate standard deviation instead of s.e.m, which are too small to display (n= 11866 – 26313 cells).

We also assessed the overall integrity of the endolysosomal pathway. For this, we incubated WT and HID-1 KO cells with EGF labeled with Alexa647 (EGF-A647) on ice, washed unbound EGF, and monitored the fluorescence by confocal microscopy over time (**Figure 2C**). After a short chase (20 min), we observed that the majority of EGF-A647 signal was found in punctate structures near the periphery of the cells, suggesting that EGF had undergone endocytosis. We observed no difference between WT and HID-1 KO cells. After a longer chase (210 min), as expected, the fluorescence intensity had decreased and was mostly found in the perinuclear region of the cells. Again, we did not observe any striking differences between the two cell types. To better quantify EGF degradation, we repeated the same assay and measured the change in fluorescence intensity by flow cytometry, and found no difference between WT and HID-1 KO PC12 cells (**Figure 2D**). We conclude that loss of HID-1 does not dramatically perturb the endolysosomal pathway. Altogether, these results suggest that HID-1, which is preferentially expressed by specialized secretory cells, specifically influences the RSP of mammalian neuroendocrine cells.

### HID-1 contributes to LDCV biogenesis by influencing TGN acidification

Our observations suggest that HID-1 might be directly involved in LDCV biogenesis. We thus tested whether the loss of HID-1 influences the composition of LDCVs. Relying on equilibrium sedimentation through sucrose, we found a pronounced shift of SgII to lighter fractions (~1M sucrose compared to ~1.5M) in HID-1 KO cells (**Figure 3A**). Interestingly, the loss of HID-1 also redistributed the calcium sensor synaptotagmin 1 away from the WT LDCV peak (bottom of the gradient) towards lighter fractions (**Figure 3B**), but had no effect on the distribution of the synaptic like microvesicle marker synaptophysin (**Figure 3C**). The change in the steady-state distribution of both soluble and transmembrane LDCV markers indicates that HID-1 influences LDCV biochemical properties and might indeed contribute to biogenesis. To complement our cell fractionation characterization, we also analyzed the cells at the ultrastructural level by electron microscopy, and observed a substantial reduction (~70%) in the number of distinguishable LDCVs in HID-1 KO cells (**Figure 4A,B**). Of note, the few remaining LDCVs displayed a decrease in dense core size (**Figure 4C**), suggesting that the observed shift in SgII density could be attributed to a decrease in the amount of cargo per vesicle. The absence of HID-1 thus affects both the number and morphology of LDCVs.

**Figure 3.**
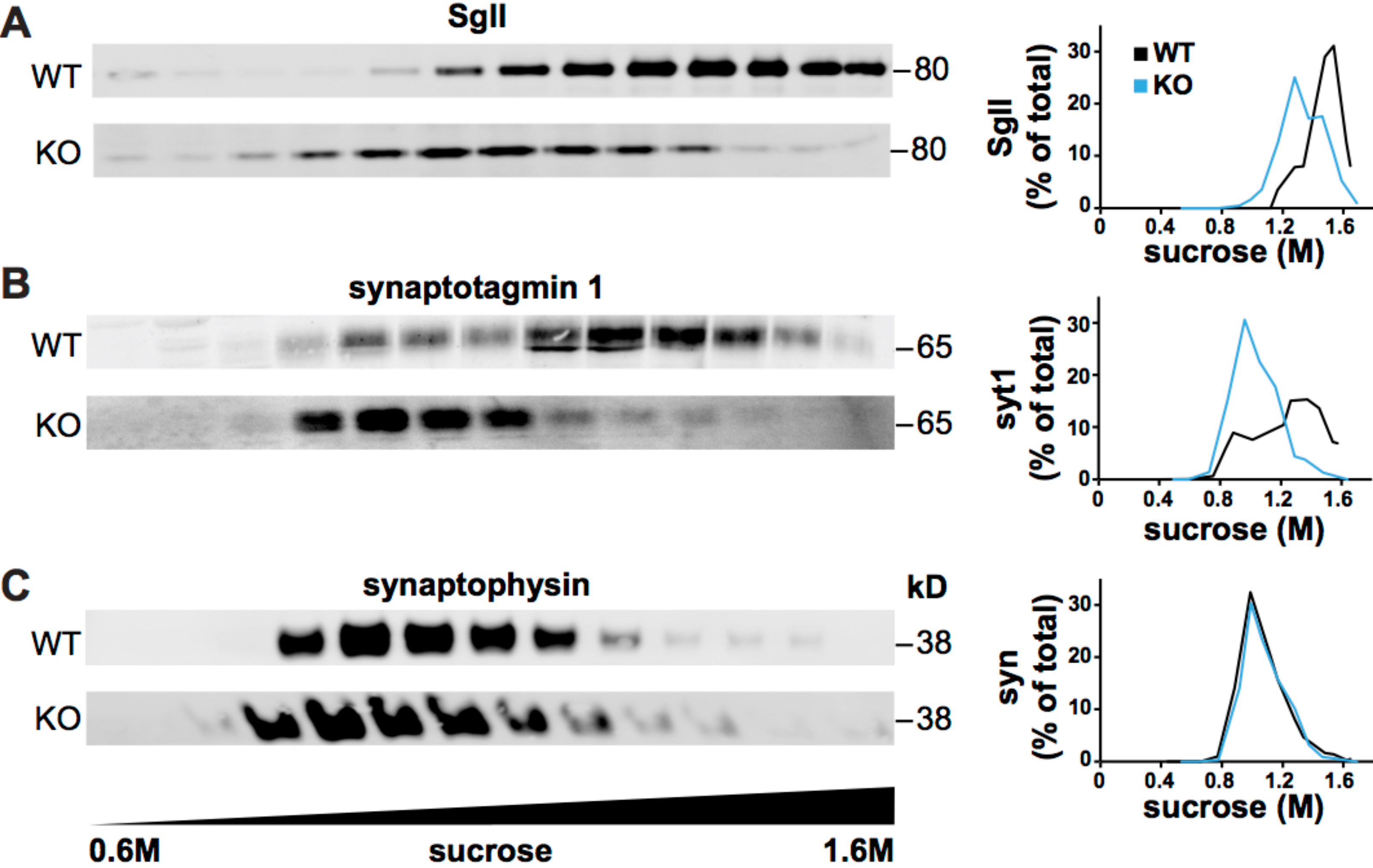
HID-1 KO cells display LDCVs with altered biochemical properties. Postnuclear supernatants obtained from WT and HID-1 KO PC12 cells were separated by equilibrium sedimentation through 0.6-1.6 M sucrose. Fractions were collected from the top of the gradient, and assayed for secretogranin II (SgII) (**A**), synaptotagmin 1 (syt1) (**B**), and synaptophysin (syn) (**C**) by quantitative fluorescent immunoblotting. The graphs (right) quantify the immunoreactivity in each fraction expressed as a percent of total gradient immunoreactivity from one experiment. Similar results were observed in an additional independent experiment.

**Figure 4.**
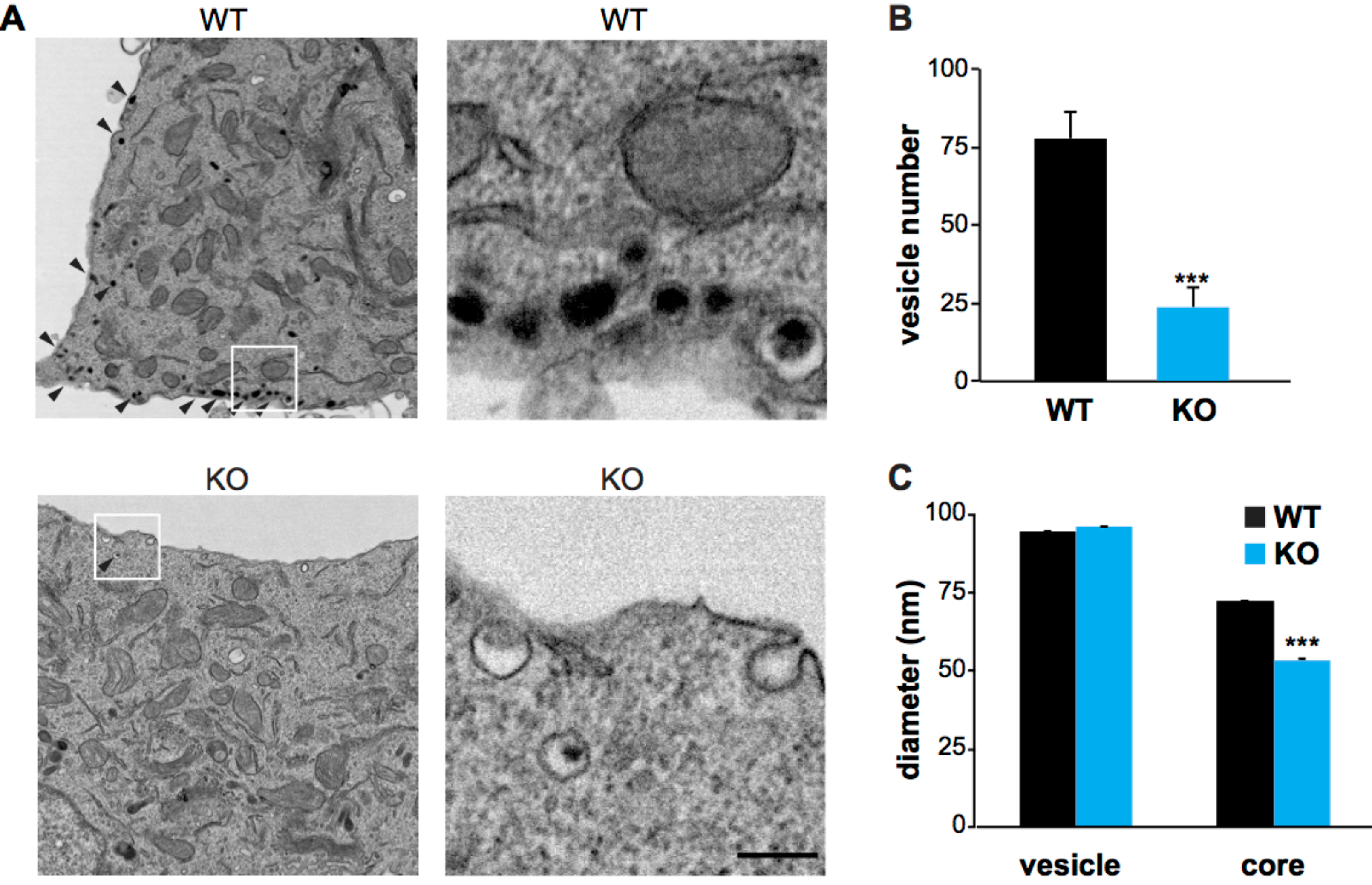
HID-1 KO cells show reduced LDCVs with abnormal morphology. (**A**) Electron micrographs (left) show a large reduction in the number of LDCVs (black arrowheads) in HID-1 KO PC12 cells relative to controls. Zoomed in regions (denoted with a white square) are shown on the right and illustrate that the absence of HID-1 leads to LDCVs with no core or dramatically reduced dense cores. The scale bar indicates 200 nm. Bar graphs indicate the number of LDCVs per cell section (n=25 cells/WT and n=21 cells/KO) (**B**) and the diameter of the vesicles and cores (n=1944 LDCVs/WT and n= 498 LDCVs/KO). ***, p < 0.001 relative to WT.

The presence of LDCVs with smaller and lighter dense cores could indicate a defect in aggregation of granulogenic proteins such as the granins. The acidic environment of the TGN lumen is thought to drive this aggregation in PC12 cells (Chanat and Huttner, 1991). As HID-1 colocalizes with Golgi markers (Wang et al., 2011), we hypothesized that HID-1 might contribute to TGN acidification. To test this directly, we targeted pHluorin to the lumen of the TGN using the transmembrane domain of the TGN marker sialyltransferase (Wong et al., 1992). We confirmed that our reporter indeed localizes to the TGN by transfecting WT and HID-1 KO PC12 cells with TGN-pHluorin and comparing its steady state localization to TGN38 by immunofluorescence (**Figure 5A**). Next, we used live-imaging to measure the fluorescence of our reporter expressed in WT and HID-1 KO PC12 cells under basal conditions. We incubated our cells with nigericin/monensin (Wu et al., 2000) to generate individual pH calibration curves for every cell that we imaged (**Figure 5B**). From the sigmoidal curves, we extrapolated absolute TGN pH values for WT and HID-1 KO cells. Strikingly, we found that the absence of HID-1 led to a significant alkalinization of the TGN (**Figure 5C**). Importantly, this phenotype could be rescued by transient expression of HID-1-HA.

**Figure 5.**
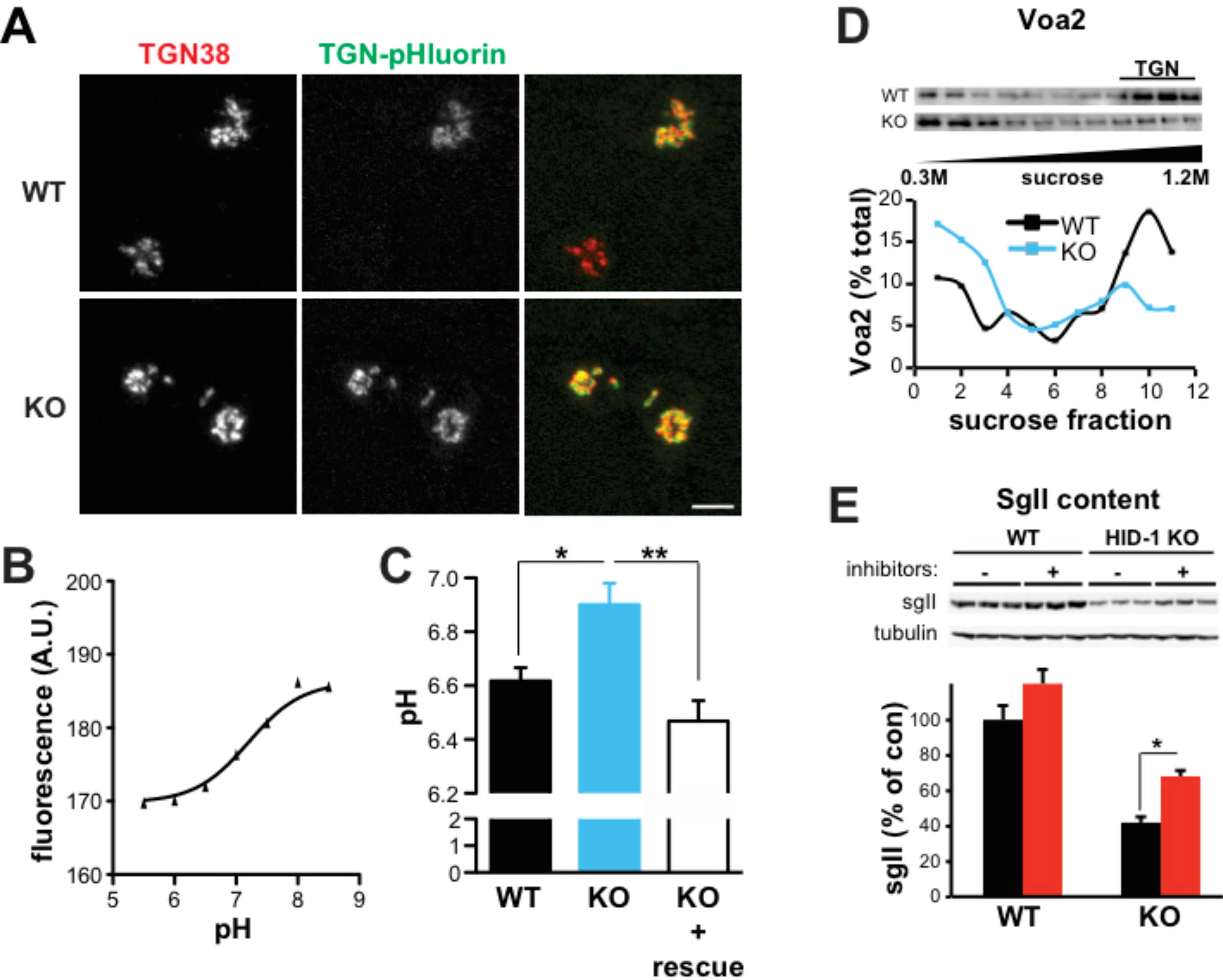
HID-1 is required for TGN acidification and soluble cargo sorting to LDCVs. (**A**) WT and HID-1 KO PC12 cells were transfected with TGN-pHluorin (green), fixed and stained with a mouse monoclonal antibody to TGN38, followed by an anti-mouse antibody conjugated to Alexa Fluor 647 (red). The scale bar indicates 5 μm. (B,C) WT and HID-1 KO PC12 cells were transfected with TGN-pHluorin and with HID-1-HA where indicated (rescue), incubated for 72 hours and imaged under basal conditions. For each cell, an individual calibration curve (**B**) was obtained by perfusing solutions of decreasing pH (8.5 to 5.5) in presence of nigericin and monensin (see Material and Methods) and used to extrapolate absolute pH values (**C**). Statistical analysis of pH values was performed by ANOVA followed by posthoc Tukey test *, p < 0.05; **, p < 0.01 relative to KO; (n=13-17 cells). The data shown indicate mean ± s.e.m. (**D**) Postnuclear supernatants obtained from WT and HID-1 KO PC12 cells were separated by velocity sedimentation through 0.3-1.2 M sucrose. Fractions were collected from the top of the gradient, and assayed for Voa2 by immunoblotting. The graph (bottom) quantifies the immunoreactivity in each fraction expressed as a percent of total gradient immunoreactivity from one experiment. Similar results were observed in an additional independent experiment. (**E**) WT and HID-1 KO PC12 cells were incubated where indicated with protease inhibitors (P.I.) for 24 hours. Cellular SgII was quantified by fluorescent western analysis and the values normalized to tubulin immunoreactivity. *, p < 0.05 relative to KO; (n=3). The data shown indicate mean ± s.e.m.

How does a peripheral membrane protein such as HID-1 influence TGN pH? The acidification of intracellular organelles depends primarily on the activity of the vacuolar H^+^-ATPase (V-ATPase), which consists of two multisubunit sectors (V_1_ and V_0_). The cytoplasmic V1 sector mediates ATP hydrolysis, whereas V_0_ assembles into a transmembrane pore involved in proton translocation (Cipriano et al., 2008, Forgac, 2007, Marshansky and Futai, 2008). PC12 cells express three isoforms (a1, a2 and a3) of the largest V_0_ subunit, and each isoform displays a particular subcellular localization with enrichment at specific organelles (Saw et al., 2011). As a2 accumulates at the Golgi in PC12 cells, this isoform is likely to contribute significantly to Golgi acidification. Thus, in order to test whether V-ATPases contribute to the phenotype caused by the lack of HID-1, we looked at the distribution of a2. For this, we relied on velocity sedimentation of PC12 membranes through sucrose to separate organelles based on their size with the Golgi migrating at the bottom of the gradient under these conditions (Asensio et al., 2010, Tooze and Huttner, 1990). Using WT membranes, a2 displayed a bi-modal distribution with a major peak at the bottom of the gradient, consistent with its reported Golgi localization (**Figure 5D**). Strikingly, the loss of HID-1 led to a dramatic redistribution of a2 to the top of the gradient away from Golgi fractions (**Figure 5D**). We thus conclude that HID-1 contributes to TGN acidification by controlling the proper accumulation and/or retention of a2 at the Golgi.

LDCV soluble cargo aggregation is thought to be a key determinant for efficient sorting to the RSP. Indeed, treatment with drugs neutralizing intraluminal pH redirects adrenal corticotropin hormone or granins to the constitutive secretory pathway (Gerdes et al., 1989, Moore et al., 1983, Taupenot et al., 2005). Surprisingly, neither our biochemical secretion experiments nor our live-imaging experiments with NPY-pHluorin (**Figure 1**) revealed any increase in basal exocytosis, which would be expected from cargo being rerouted to the constitutive secretory pathway. Then, where does the soluble cargo go? Previous work has shown that reduced levels of LDCV soluble cargo observed in *hid-1* mutant worms could be partially rescued by genetically inhibiting lysosome biogenesis (Yu et al., 2011). To test whether increased lysosomal degradation is contributing to the depletion of SgII observed in HID-1 KO PC12 cells, we incubated our cells with protease inhibitors for 24 hours to inhibit lysosomal hydrolases (Asensio et al., 2013, Sirkis et al., 2013), and found that the treatment partially rescued SgII levels in HID-1 KO cells, suggesting that a portion of SgII is redirected to the lysosome in absence of HID-1 (**Figure 5E**).

### HID-1 influences the sorting of transmembrane LDCV cargo

Our data indicate that HID-1 influences sorting of soluble LDCV cargo by impairing their aggregation into a dense core, but does it contribute to sorting of transmembrane LDCV proteins as well? Similar to chromaffin cells, PC12 cells store monoamines into LDCVs through the action of an integral membrane protein: the vesicular monoamine transporter (VMAT), which depends on a cytoplasmic di-leucine-like motif for sorting to LDCVs (Erickson et al., 1995; Liu et al., 1994). Mutations within this motif cause an increase in VMAT cell surface delivery by diverting the transporter from the regulated to the constitutive secretory pathway (Asensio et al., 2010, Li et al., 2005). To test whether HID-1 influences sorting of VMAT, we monitored its cell surface delivery using a flow cytometry assay that we have previously developed (Asensio et al., 2010). For this, we transfected WT and HID-1 KO PC12 cells with HA-VMAT2-GFP, which contains a lumenal HA tag to monitor surface level as well as a cytosolic GFP to determine total expression. After incubating the transfected cells with an anti-HA antibody conjugated to Alexa647, we measured GFP and Alexa647 fluorescence by flow cytometry. Cumulative frequency distribution of the ratio of surface (HA) over total (GFP) showed no difference between WT and HID-1 KO PC12 cells. As a positive control, we observed that WT cells transfected with EE/AA HA-VMAT2-GFP, a trafficking mutant that missorts to the plasma membrane (Asensio et al., 2010, Li et al., 2005), exhibited the expected shift in distribution (**Figure 6A**). VMAT2 is thus not being rerouted to the constitutive secretory pathway by default in absence of HID-1.

**Figure 6.**
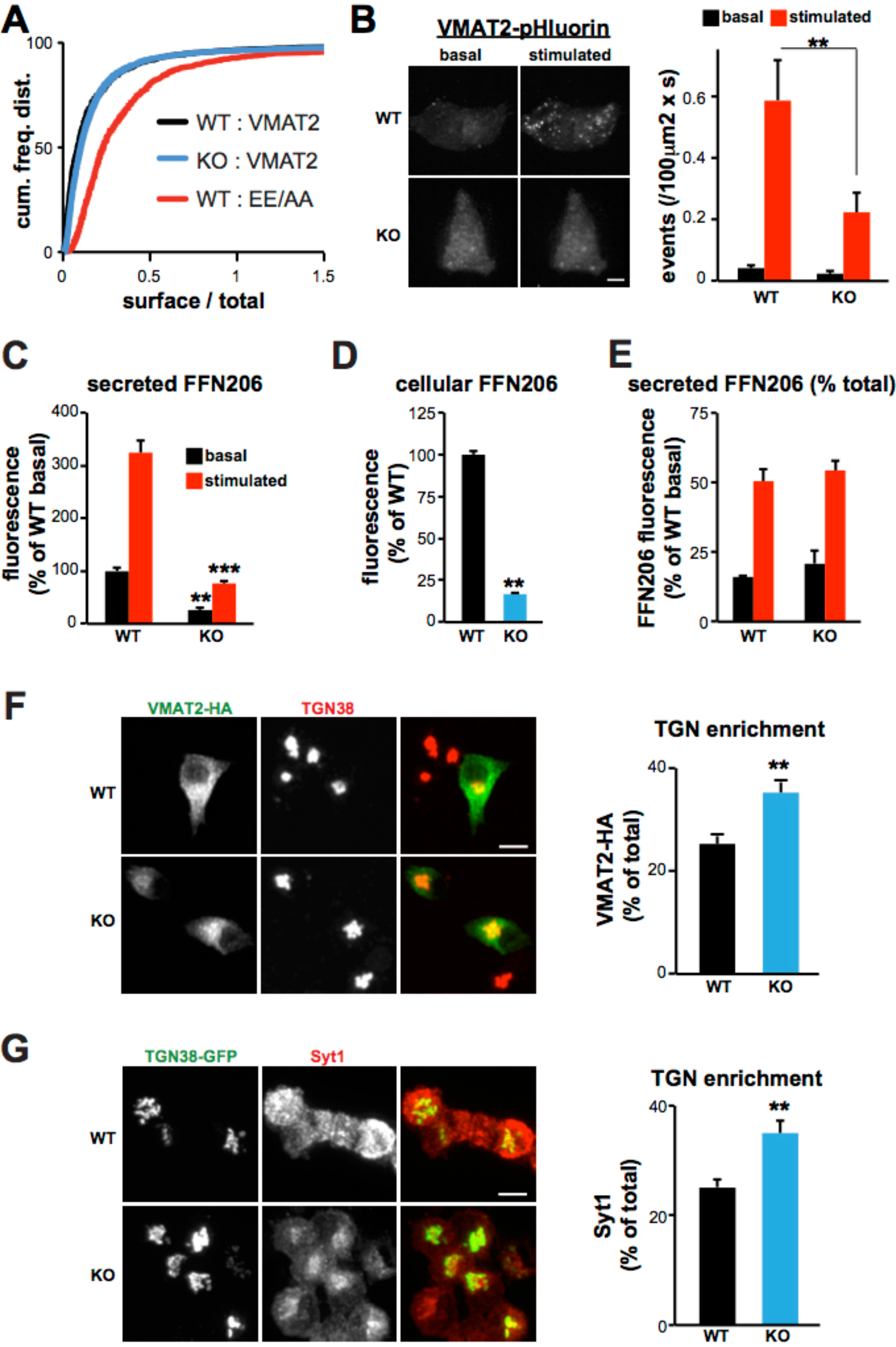
Loss of HID-1 leads to TGN enrichment of transmembrane LDCV cargoes. (**A**) WT and HID-1 KO PC12 cells were transfected with the indicated HA-VMAT2-GFP constructs, incubated with an HA antibody conjugated to Alexa Fluor 647 for 1 h, washed and analyzed by flow cytometry to determine the fluorescence of individual cells. The ratios of surface (HA) over total (GFP) were computed and expressed as a cumulative frequency distribution. (**B**) WT and HID-1 KO PC12 cells were transfected with VMAT2-pHluorin, then imaged live by spinning disk confocal microscopy as described in **Figure 1**. Images show representative maximum intensity time projections of 150 basal and stimulated frames. At the end of the experiment, cells were imaged in Tyrode’s solution containing 50 mM NH_4_Cl, pH 7.4 to reveal total VMAT2-pHluorin fluorescence by alkalinization and identify transfected cells. Scale bar indicates 5 μm. Bar graph shows the number of exocytotic events per second normalized to cell surface area. Statistical analysis was performed by one-way ANOVA followed by posthoc Tukey test. **, p < 0.01 relative to stimulated secretion from WT (n=10-14 cells from two independent experiments). The data shown indicate mean ± s.e.m. (**C-E**) PC12 cells were loaded with FFN206 (a vesicular monoamine transporter fluorescent substrate) and subjected to a secretion assay as described in Fig 1. Secreted (**C**) and cellular (**D**) fluorescence values were measured using a plate reader. (**E**) Secreted FFN206 was expressed as percent of total FFN206 fluorescence. Statistical analysis was performed by one-way ANOVA followed by posthoc Tukey test. **, p < 0.01 relative to basal secretion from WT; ***, p < 0.001 relative to stimulated secretion from WT (n=8). The bar graphs indicate mean ± s.e.m. (**F**) WT and HID-1 KO PC12 cells were transfected with VMAT2-HA, fixed and stained with HA and TGN38 antibodies followed by Alexa Fluor 488- and Alexa Fluor 647-conjugated secondary antibodies or (**G**) WT and HID-1 KO PC12 cells were transfected with TGN38-GFP, fixed and stained with Syt1 antibody followed by Alexa Fluor 647-conjugated secondary antibody. The cells were imaged by spinning-disk confocal microscopy. The amount of HA (**F**) or Syt1 (**G**) immunoreactivity overlapping with TGN38 was expressed as a percent of total fluorescence. **, p < 0.001 relative to WT (n=16-30 cells). The scale bars indicate 5μm. The data shown indicate mean ± s.e.m.

This observation raises the intriguing possibility that VMAT2 might be able to sort onto LDCVs lacking a dense core. To test this directly, we transfected WT and HID-1 KO PC12 cells with VMAT2 fused to pHluorin (Asensio et al., 2010, Onoa et al., 2010) and monitored individual exocytotic events using live-imaging (**Figure 6B, Movies S3,4**). As expected, WT PC12 cells displayed very few events under basal conditions, but showed a massive response to depolarization. Consistent with our flow cytometry data, the number of non-stimulated exocytotic events in HID-1 KO cells remained unchanged, again suggesting that VMAT2 is not being rerouted to the constitutive secretory pathway. However, we observed a dramatic reduction in the amount of events triggered by depolarization, suggesting that VMAT2 does not sort onto “empty” LDCVs in absence of HID-1. As a complementary approach, we developed an assay to assess monoamine secretion taking advantage of the false fluorescent neurotransmitter (FFN206), which is a VMAT substrate (Hu et al., 2013). After loading WT and HID-1 KO PC12 cells with FFN206, we measured the fluorescence of secreted and cellular fractions under basal and stimulated conditions using a plate reader. Similar to the effect observed with SgII, we found that loss of HID-1 led to a decrease in basal storage as well as both basal and regulated secretion of FFN206. However, normalization of the secretion data to the cellular content showed no difference in regulated secretion *per se* (**Figure 6C-E**). Altogether, these data indicate that the absence of HID-1 reduces the number of VMAT2-positive LDCVs available for regulated exocytosis, but also suggest that VMAT2 does not sort onto “empty” LDCVs.

If the number of VMAT2 positive LDCVs available for release is reduced, but VMAT2 does not traffic to the plasma membrane constitutively, where does excess VMAT2 accumulate? To address this, we immunostained WT and HID-1 KO PC12 cells transfected with VMAT2-HA, and determined the steady-state localization of the transporter. The images revealed VMAT2 enrichment in the perinuclear area, suggesting that the transporter might get retained in the Golgi area. Consistent with this, we observed an increase in the amount of VMAT2 overlapping with TGN38 (**Figure 6F)**. Finally, we also determined the steady-state localization of an endogenous transmembrane LDCV cargo. Immunostaining for Syt1 showed that this marker similarly accumulates in the TGN area (**Figure 6G**).

## Discussion

These results establish that the peripheral membrane protein HID-1 is required for LDCV biogenesis from rat neuroendocrine PC12 cells. The absence of HID-1 leads to defects in both storage and secretion of SgII as well as monoamines. Our phenotype is consistent with earlier studies in worms showing a decrease in accumulation and release of neuropeptides (Mesa et al., 2011, Yu et al., 2011). More recently, it has been reported that conditional KO mice, in which HID-1 has been specifically inactivated from beta-cells of the pancreas, display impaired insulin secretion. This phenotype has been attributed to a defect in homotypic fusion of immature insulin granules (Du et al., 2016), a mechanism that is somewhat surprising considering that this fusion process had never been previously observed in this cell type. Although we cannot definitively rule out that this might contribute to our phenotype, we did not observe any obvious defects in LDCV homotypic fusion in our HID-1 KO PC12 cells by electron microscopy. It is also important to note that immature LDCVs can still undergo exocytosis in PC12 cells (Tooze et al.,1991). Thus, it is unlikely that our secretion defect is caused uniquely by a maturation defect. Instead, given its enrichment at the Golgi, we propose that HID-1 controls a step upstream of LDCV maturation, probably at the level of budding at the TGN.

In agreement with this, we found that the absence of HID-1 leads to a striking mislocalization of a2 concomitant with a significant alkalinization of the TGN. This defect in TGN acidification is likely to prevent dense core formation by impairing granin aggregation. Indeed, we observed a dramatic reduction in the total number of vesicles with a discernable dense core by electron microscopy. In addition, the few remaining LDCVs exhibited smaller cores and displayed abnormal biochemical properties with a shift of SgII towards lighter sucrose fractions after equilibrium sedimentation. Given the known significance of core formation for efficient sorting of soluble cargo to the RSP, we expected SgII to be missorted to the constitutive secretory pathway by default. Surprisingly, basal secretion of SgII remained unchanged in HID-1 KO cells, and instead, we found that a portion of SgII was rerouted to the lysosome. It is important to note that drugs that have been used in the past to neutralize TGN pH, which led to rerouting of soluble LDCV cargo to the constitutive secretory pathway (Gerdes et al., 1989, Moore et al., 1983, Taupenot et al., 2005), also impair lysosome function. Thus missorting of cargo to the lysosome might have previously been overlooked. We propose that lack of SgII aggregation leads to its dilution and diffusion throughout the endomembrane system (**Figure 7**). Alternatively, defective LDCVs might be targeted for degradation after budding from the TGN by crinophagy (Orci et al., 1984).

**Figure 7.**
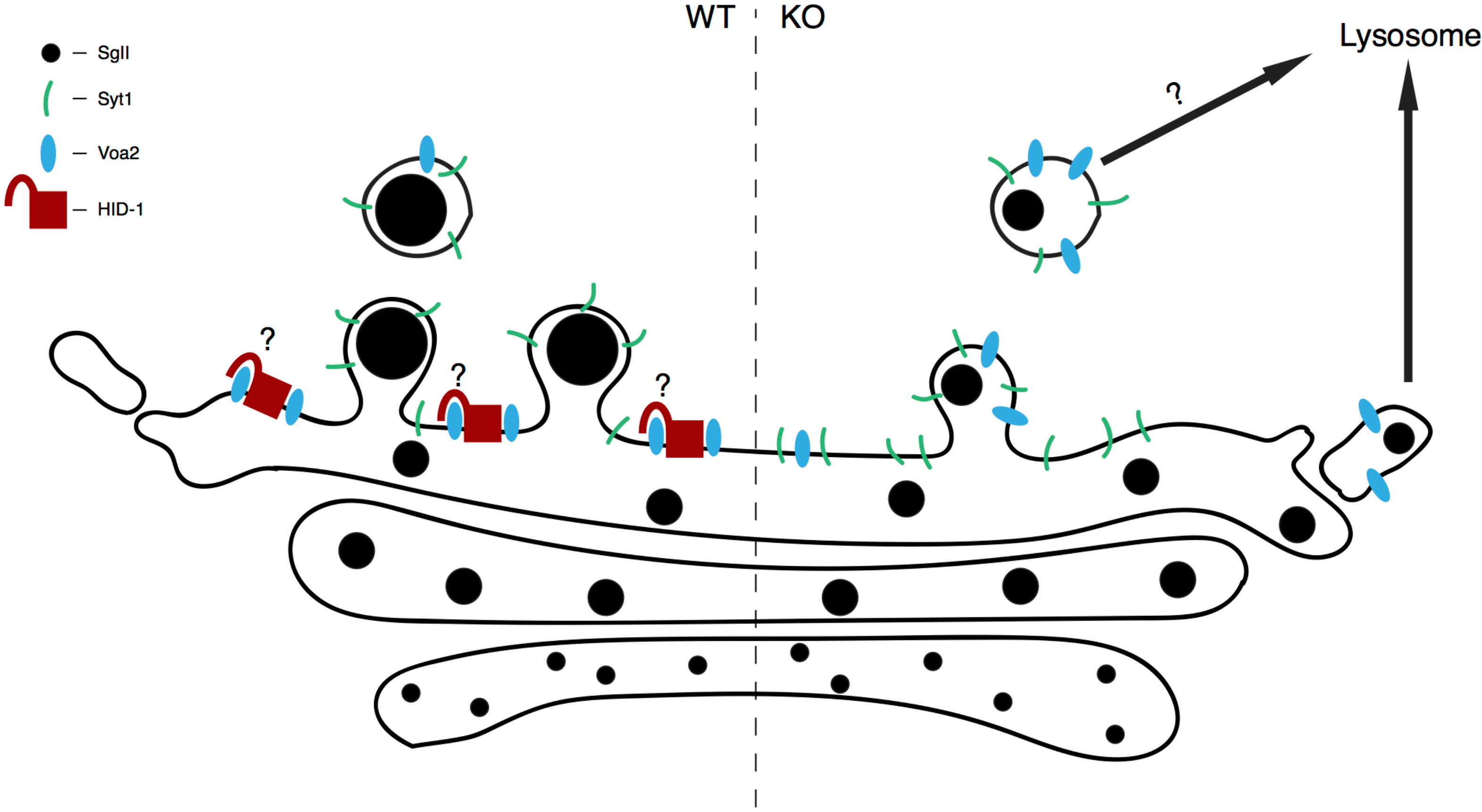
Proposed model for the function of HID-1 at the TGN. HID-1 acts as a gatekeeper preventing Voa2 from diffusing out of the TGN. Loss of HID-1 prevents TGN acidification, which in turn impairs LDCV soluble cargo condensation and dense core formation. Disruption of protein aggregation leads to diffusion of soluble LDCV cargos throughout the endomembrane system with a portion being routed to the lysosome for degradation. Inefficient core formation effectively reduces the number of budding LDCVs and causes LDCV-resident transmembrane proteins to accumulate at the TGN. Alternatively, defective LDCVs could be degraded by crinophagy.

As the expression of HID-1 is restricted to specialized secretory cells (Wang et al., 2011), it raises the question of why would HID-1 be needed in some cell types but not others. Indeed, the existence of an acidic Golgi apparatus is not specific to secretory cells but rather a general feature of eukaryotic cells. It is tempting to speculate that HID-1 acts as a gatekeeper to prevent a2 from leaking into the RSP (**Figure 7**). In PC12 cells, knockdown of a1, the isoform enriched on LDCVs, is not sufficient to impair LDCV acidification, but double knockdown of both a1 and a2 leads to alkalinized LDCVs, suggesting that a2 has the propensity to escape onto LDCVs (Saw et al., 2011). In absence of an efficient retention mechanism, the continuous budding of LDCVs and other vesicles might effectively deplete a2 from the TGN. It will be interesting to test this in the future by assessing whether HID-1 is able to interact directly with a2 or whether it mediates its effect indirectly through another partner. Another explanation could be related to the biochemical properties of the secretory cargo that specialized secretory cells need to accommodate. Indeed, with a predicted isoelectric point of ~4.7, both SgII and chromogranin A are acidic proteins carrying a net negative charge at neutral pH. As these proteins are expressed at very high level in most specialized secretory cells, they might significantly increase the buffering capacity of the biosynthetic pathway of these cells making it harder to build a pH gradient. These secretory cells might thus rely on some additional mechanisms to ensure efficient Golgi acidification and HID-1 might play an important role by influencing a2 localization.

We have previously observed that knockdown of AP-3 or VPS41 leads to defects in SgII storage and regulated secretion together with constitutive delivery of transmembrane LDCV cargoes to the cell surface suggesting that these cytosolic factors recruit and concentrate membrane proteins onto LDCVs (Asensio et al., 2010, Asensio et al., 2013). In striking contrast, transmembrane LDCV cargoes do not traffic constitutively to the plasma membrane in absence of HID-1, but instead, accumulate in a perinuclear compartment where they partially overlap with TGN markers. In addition, although loss of HID-1 also results in defects in SgII storage, regulated secretion *per se*, when normalized to cellular stores, remains unchanged. Thus, the few LDCVs budding from the TGN of HID-1 KO cells must be fully competent for regulated exocytosis, and presumably have the proper membrane protein composition. This suggests that HID-1 does not influence the ability of the cytosolic sorting machinery to interact with transmembrane cargoes. So, why do Syt1 and VMAT2 display TGN enrichment? We propose that inefficient core formation not only reduces the efficiency of soluble cargo sorting, but also influences the total number of budding vesicles. If transmembrane cargoes can still interact with the cytosolic sorting machinery, then reductions in the amount of budding would result in their accumulation at the TGN with the cytosolic sorting machinery effectively preventing them from leaking into the constitutive secretory pathway.

## Materials and Methods

### Molecular biology

The human codon-optimized Cas9 and chimeric guide RNA expression plasmid (pX330) developed by the Zhang lab (Cong et al. Science. 2013, PMID: 23287718) were obtained from Addgene. To generate gRNA plasmids, a pair of annealed oligos (20 bp) was ligated into the single guide RNA scaffold of pX330. The following gRNAs sequences were used: Forward1: 5′-CACCGAGCCACCGACAATGCTTTCT-3′; Reverse1: 5′-AAACAGAAAGCATTGTCGGTGGCTC-3′; Forward2: 5′-CACCGTGGCTTCCACGGGCTGCAG-3′; Reverse2: 5′-AAACCTGCAGCCCGTGGAAGCCAC-3′. For the homologous recombination repair template, tdTomato-polyA was inserted into pBSKII(+), and HID-1 homology arms were amplified by PCR and inserted 5’ and 3’ of it. The following primers were used to amplify the homologous arms: 5’ arm Forward: 5’-TCCCTGAAAGTGACTGGAGC-3’; 5’ arm Reverse: 5’-CTGCAGGGGAATCGAGTAGC-3’; 3’ arm Forward: 5’-CTTTCTGGGACCAGTTCTGG-3’;3’ arm Reverse: 5’-TGGTATCCCTCTCTTAAGGAGTGC-3’. ssGFP was constructed by amplifying the signal sequence of rat ANF and fusing it directly to GFP into the chicken actin-based vector pCAGGs. HID-1 lentiviral plasmid was generated by amplifying HID-1 from rat PC12 cells cDNA using the following primers: WT Forward: 5’-ATGGGGTCTGCAGACTCC-3’; WT Reverse: 5’-CACCCGCTGGATCTCAAAGAG-3’. The main isoform amplified from these cells correspond to the isoform #2, which has been reported for mouse (Uniprot: Q8R1F6-2) and for human (Uniprot: Q8IV36-2). The PCR products were then subcloned into FUGW.

### Cell culture and Lentivirus Production

PC12 cells were maintained in DMEM medium supplemented with 10% horse serum and 5% calf serum in 5% CO2 at 37 °C. Transfection of PC12 cells was performed using Lipofectamine 2000 (LifeTechnologies) or Fugene HD (Promega) according to the manufacturer’s instructions. HEK293T cells were maintained in DMEM medium with 10% fetal bovine serum (FBS) in 5% CO_2_ at 37 °C. Lentivirus was produced by transfecting HEK293T cells with FUGW, psPAX2 and pVSVG using Fugene HD according to the manufacturer’s instructions.

### Antibodies

The HA.ll mouse monoclonal antibody was obtained from Covance (USA), the HA (3F10) rat monoclonal antibody from Roche, the SgII rabbit antibody from Meridian Life Science (USA), TGN38 mouse monoclonal from BD Biosciences (USA), the tubulin mouse monoclonal antibody from Developmental Hybridoma Bank Studies (USA), the synaptotagmin 1 mouse antibody from Synaptic Systems (Germany), the synaptophysin (p38) monoclonal antibody from Covance (USA), the HID-1 mouse monoclonal antibody from Novus Biologicals (USA), the goat anti-rabbitAlexa Fluor 647, anti-mouse Alexa Fluor 488, anti-mouse Alexa Fluor 647, and anti-rat Alexa Fluor 488 secondary antibodies from Molecular Probes (USA). The a2 rabbit polyclonal antibody was a generous gift from Dr. Xiao-Song Xie (UT Southwestern).

### Secretion assays

PC12 cells were plated on poly-l-lysine, washed and incubated in Tyrode’s buffer containing 2.5 mM K^+^ (basal) or 90 mM K^+^ (stimulated) for 30 minutes at 37° C. The supernatant was then collected, cell lysates prepared as previously described (Asensio et al., 2010), and the samples analyzed by quantitative, fluorescent immunoblotting. For FFN206 assays, PC12 cells were preloaded with (1 μM) FFN206 in Tyrode’s buffer for 45min. Following incubation, cells were washed twice with Tyrode’s buffer and subjected to a secretion assay as described above. Supernatant and lysates were loaded on 96well plates and fluorescence was read using a plate reader (excitation: 369nm; emission : 464nm).

### Immunofluorescence and confocal microscopy

PC12 cells were rinsed with PBS and fixed in 4% paraformaldehyde in PBS and incubated for 20 min at room temperature. Cells were permeabilized in PBS containing 0.1% Triton-X100 for 10min at RT and blocked in PBS containing 2% BSA, 1% fish skin gelatin and 0.02% saponin. Primary antibodies were diluted in blocking solution at 1:1000 (SgII), 1:1000 (TGN38), 1:1000 (Syt1), 1:500 (HA) and 1:500 (HA.11). The secondary goat anti-rabbit antibodies were diluted in blocking solution at 1:1000. Images were acquired using an Olympus Fluoview scanning confocal microscope and 63x oil objective (NA 1.42) at a resolution of 512x512 pixels with a sampling speed of 12.5μs/pixel with Kalman filter (integration count 5).

### Density gradient fractionation

Equilibrium sedimentation through sucrose was performed as previously described (Asensio et al., 2010, Asensio et al., 2013). Briefly, a postnuclear supernatant was prepared from PC12 cells by homogenization with a ball bearing device (12 µm clearance), loaded onto a 0.6 M-1.6 M continuous sucrose gradient, and sedimented at 30,000 rpm in an SW41 rotor for 14-16 h at 4 °C. Fractions (~750 µl each) were collected from the top and were analyzed by quantitative, fluorescent immunoblotting using an FX Imager (BioRad), or a FluoChem R (ProteinSimple). Velocity sedimentation through sucrose was performed as described above with postnuclear supernatant loaded onto a 0.3 - 1.2M continuous sucrose gradient, and sedimented at 25,000rpm for 19min. Fractions (1ml) were collected from the top and analyzed as above.

### Electron microscopy

PC12 cells were plated onto aclar film discs coated with poly-L-lysine, fixed with 2.5% glutaraldehyde, 2% paraformaldehyde, and 2mM calcium chloride in 0.15M cacodylate buffer (pH 7.4). Coverslips were rinsed and fixed in a mixture of 2.5% glutaraldehyde and 2% paraformaldehyde in 0.15M cacodylate buffer at a pH 7.4 with 2 mM calcium chloride warmed to 37°C for 5 minutes in an incubator. Subsequently, samples were transferred to a refrigerator at 4 °C in the same fixative solution until they were ready to be processed. Once ready to be processed, each coverslip was rinsed in 0.15 M cacodylate buffer 3 times for 10 minutes each, and subjected to a secondary fixation step for one hour in 1% osmium tetroxide/0.3% potassium ferrocyanide in cacodylate buffer on ice. Following this, samples were then washed in ultrapure water 3 times for 10 minutes each and en bloc stained for 1 hour with 2% aqueous uranyl acetate. After staining was complete, samples were briefly washed in ultrapure water, dehydrated in a graded acetone series (50%, 70%, 90%, 100%, 100%) for 10 minutes in each step, infiltrated with microwave assistance (Pelco BioWave Pro, Redding, CA) into LX112 resin, and flat embedded between two slides that had been coated with PTFE release agent (Miller-Stephenson #MS-143XD, Danbury, CT). Samples were cured in an oven at 60 °C for 48 hours. Once resin was cured, the slides were separated and the aclar coverslips were peeled off. A small region was excised and glued onto a blank stub with epoxy. 70nm thin sections were taken and imaged on a FE-SEM (Zeiss Crossbeam 540, Oberkochen, Germany) using the aSTEM detector. The SEM was operated at 28 KeV and a probe current of 0.9 nA, and the STEM detector was operated with the annular rings inverted for additional image contrast. Whole cells were imaged at 6114 x 4608 pixels with a resolution of 3.722 nm/pixel from random sections. Zoomed regions were imaged at 2048 x 1536 pixels with a resolution of 1.861 nm/pixel. Images were analyzed with ImageJ. Morphologically identifiable LDCVs were counted per cell section and performed blind to the conditions of the experiments.

### Lysosomal inhibition

PC12 cells were incubated for 24 hrs in complete medium supplemented with vehicle or a cocktail of lysosomal protease inhibitors (Sigma) including (in μM) 10 antipain, 10 leupeptin and 5 pepstatin A. Cells were washed on ice with cold PBS and lysed by the addition of 50 mM Tris-HCl, pH 8.0, 150 mM NaCl, 1% Triton X-100, and SIGMAFAST Protease Inhibitor Cocktail (Sigma-Aldrich) plus 10 mM EDTA and 1 mM PMSF. Samples were analyzed by quantitative fluorescent immunoblotting.

### EGF degradation assay

WT and HID-1 KO PC12 cells were washed twice with PBS and starved of serum for 2hrs in DMEM with 0.1% BSA (GoldBio). During starvation, EGF-biotin (GoldBio) streptavidin-647 conjugate was prepared. EGF-biotin (5ug/mL) was incubated for 30min at 4 °C at a 5:1 ratio to streptavidin-Alexa647 (Life Technologies). Following starvation, cells were washed twice with ice cold PBS on ice and incubated with EGF-A647 conjugate at a final concentration of 100ng/mL for 1 hr on ice. Excess unbound EGF was removed by washing with ice cold PBS with 0.5% BSA. Cells were chased for indicated times before fixation with 4% PFA in PBS for 20min at room temperature. Cells were analyzed by flow cytometry (CyAn ADP Analyzer, Beckman Coulter, USA) or by scanning confocal microscopy.

### Spinning disk confocal live-imaging

PC12 cells were co-transfected with NPY-pHluorin or VMAT2-pHluorin together with BDNF-mCherry. One day after transfection, cells were transferred to poly-L-lysine coated 22 mm glass coverslips. After an additional two days, cells were washed once with Tyrode’s buffer and coverslips were transferred to an open imaging chamber (Life Technologies). Cells were imaged close to the coverslips focusing on the plasma membrane (determined by the presence of BDNF-mCherry positive plasma membrane docked-vesicles) using a custom built Nikon spinning disc at a resolution of 512x512 pixels. Images were collected for 100 ms at 10 Hz at room temperature with a 63x objective (Oil Plan Apo NA 1.49) and an ImageEM X2 EM-CCD camera (Hamamatsu, Japan). Following baseline data collection (15 s), an equal volume of Tyrode’s buffer containing 90mM KCl was added to stimulate secretion and cells were imaged for an additional 30 s. At the end of the experiment, cells were incubated with Tyrode’s solution containing 50 mM NH_4_Cl, pH 7.4 to reveal total fluorescence and to confirm that the imaged cells were indeed transfected. Movies were acquired in MicroManager (UCSF), and exported as tif files. To automatically detect newly-appearing exocytic events within a cell, difference images were constructed between the averages of adjacent pairs of frames in a given movie, i.e. mean (n+2, n+3)-mean (n, n+1) where n is any frame between the first and last frame minus three. Positive differences in intensity between the averaged frames were taken to be candidate events. The difference images were then passed through a Gaussian filter to reduce image noise. Transfected cells were cropped from each movie by hand and analyzed individually. Event detection was performed using a wavelet-based method as previously described (Jaqaman et al., 2008, Olivo-Marin, 2002). Noise events were filtered from the true events on the basis of intensity and the stipulation that events occur within the confines of the cell, and not overlap with previously counted events. A convex hull around the remaining events was generated to approximate cell area and to find the density of events. Cell activity could then be assessed by producing the cumulative sum of events produced by a cell over the course of the movie. All image analysis was performed using MATLAB and the MATLAB Image Processing Toolbox.

### pH imaging

PC12 cells were transfected with TGN-pHluorin. One day after transfection, cells were transferred to poly-L-lysine coated 22mm glass coverslips. After an additional two days, cells were washed once with Tyrode’s buffer (pH 7.4) and imaged using a Zeiss Axiovert 5100TV widefield microscope. Images were collected with 100 ms exposure at a resolution of 512x512 pixels at room temperature with a 40x objective (NA 1.30) and a CoolSNAP HQ2 camera (Photometrics, USA). Cells were then perfused with enriched KCl buffer supplemented with 5μM nigericin (Sigma-Aldrich) and 5nM monensin (Sigma-Aldrich) at pH 8.5 and incubated for 10 min before image acquisition. This process was repeated in pH increments of 0.5 down to pH 5.5.

### Quantitative PCR

RNA was isolated from HID-1 KO or WT PC12 cells with the E.Z.N.A Total RNA Isolation Kit (Omega), then isolated RNA was DNAse treated with Turbo DNAse (Ambion). 2ug of total RNA was reverse transcribed (SuperScriptIV, Thermo) and subjected to triplicate qPCR for the last exon-exon junction of ActB and SgII transcripts. qPCR was performed using SYBR Green qPCR Master Mix (BioRad) and BioRad iQ5 Real-Time PCR machine (BioRad) with gene specific primers. The results were normalized to expression of the house-keeping gene ActB. SgII forward primer: CCTACTTGAGAAGGAATTTGC SgII Reverse primer: ACCAACCCATTTGGTTTCTC ActB Forward primer: CCTAGCACCATGAAGATCAA ActB Reverse primer: GATAGAGCCACCAATCCAC.

### Statistics

Unless indicated otherwise, all statistical analysis was performed using the two-tailed Student’s t-test. Statistical analyses were conducted using Excel or Prism.

### Figure preparation

Images were processed using ImageJ, any changes in brightness and contrast were identical between samples meant for comparison.

## Acknowledgments

We thank Dr. Emmanuel Boucrot and Dr. Dan Sirkis for critical reading of the manuscript, Dr. Dinah Loerke for help with image analysis, the Angleson lab for access to their microscope, members of the Asensio lab for thoughtful discussions, Josh Loomis and Shirley Sobus for help with cell sorting and analysis at the National Jewish Health Center flow cytometry core facility (Denver), Dr. Xiao-Song Xie for the a2 antibody, the National Institute of Health (GM116096), the American Heart Association (#16BGIA27780049) and the American Diabetes Association (1-17-JDF-064) for support to CSA. MSJ and JAJF gratefully acknowledge support from the Washington University Center for Cellular Imaging, which is supported by Washington University School of Medicine, the Children’s Discovery Institute of Washington University and St. Louis Children’s Hospital (CDI-CORE-2015-505) and the Foundation for Barnes Jewish Hospital.

